# Comparisons of Tree Damage Indicators in Five NASA ABoVE Forest Sites Near Fairbanks, Alaska

**DOI:** 10.1101/2024.07.10.602861

**Authors:** Diane C. Huebner, Christopher S. Potter

**Affiliations:** Oak Ridge Associated Universities; NASA Ames Research Center

**Keywords:** Alaska, boreal, browning, burn severity, herbivory, non-photosynthetic carbon, pathogens, tree damage

## Abstract

As global warming affects sensitive northern regions, forests near Fairbanks, Alaska may be undergoing attack from pests and pathogens that could impact their ability to store carbon. Visual tree surveys are quick and useful for assessing forest health in remote sensing studies using GT (ground-truthing). Initial spectral analysis of leaf pigments, canopy water content, and non-photosynthetic carbon of one site near Fairbanks, Alaska imaged with AVIRIS-NG by NASA for the Arctic and Boreal Vulnerability Experiment (ABoVE) showed high fire fuel loads in 2017 that burned in 2019. In 2021-2022 we visually assayed damage of 359 deciduous and 309 coniferous trees at five ABoVE sites of different moisture regimes and burn severities. Using indices of 0 - 5 (0 = healthy, 5 = severe damage) we calculated average damage per tree from: 1) leaf damage (holes or defoliation); 2) stem damage (changes in stem color, texture, growth, heartwood, sap ooze, or stem loss); 3) non-photosynthetic tissue, aka “browning”; and 4) wilting. We also characterized crown color tree-1. Least squares models found low overall average tree damage, but damage types were varied and complex. Deciduous trees suffered greater herbivore damage than conifers. A third of trees showed broadleaf insect damage, a tenth of trees across species showed stem damage associated with pathogens. Aspen and conifers showed heartwood rot, but we found no visual signs of spruce beetle at our sites. Structural equation models found greater stem damage and wilting in warmer soils and post-burned sites supporting seedlings. Browning was associated with understory branches of conifers in late-successional sites with colder, shallower soils. Our study suggests that deciduous trees and seedlings near Fairbanks, Alaska are experiencing herbivory and midsummer wilting, and conifer understory browning is common.

## Introduction

Climate warming is exposing the world’s northern forests to unprecedented stress, and their long-term resilience to rapid change remains unknown (Trumbore et al. 2015). Over the past half century, Alaska has warmed in excess of 0.5 °C per decade, at roughly twice the rate of the contiguous United States (Ballinger et al. 2023). 2019 was Alaska’s warmest year on record; 2023 recorded the hottest Pan-Arctic summer (Ballinger et al. 2023b). Mean annual precipitation (MAP) in Alaska’s Interior is historically variable (AICC 2024); the drying trend seen in Alaska’s Eastern Interior in the previous decade (Parent and Verbyla 2010) currently shows a wetter trend (Ballinger et al. 2023; SNAP 2024), which may offset drought stress In the Tanana Valley. Alternatively, increased net primary productivity (NPP) has been shown to accelerate soil drying in the Northern Hemisphere through increased evapotranspiration of plants (Li et al. 2016), particularly at deeper soil depths (Wei et al. 2022), which could be accelerated with increased warming. Alaskan spruce has shown reduced growth response to drought (Barber et al. 2000; Wilmking and Myers-Smith 2008; Walker et al. 2015) that may be magnified by pests and pathogens, leading to increased boreal browning (Parent and Verbyla 2010). Warming temperatures could also benefit early larval development of insect defoliators such as aspen leaf miner (*Phyllocnistis populiella* (Chambers)), at least in the short term (Woods et al. 2022), which could extend the season of attack.

Since the early 20^th^ century, the US Department of Agriculture (USDA) Forest Service has conducted forest health surveys in the contiguous United States and Alaska as part of the Forest Inventory and Analysis (FIA) Program (Smith 2002), albeit spatial and temporal gaps exist in survey records for Alaska (Potter and Conkling 2017; Randolph et al. 2021). Since 2002, *P. populiella* outbreaks have been recorded in the Alaskan Interior, reaching 300,000 ha of Alaskan forest by 2008 (Wagner and Doak 2013). Since 2016, spruce beetle (*Dendroctonus rufipennis* (Kirby)) has affected nearly 2 million acres of forest in South-Central Alaska, leading to needle loss and mortality of coniferous species, mainly white spruce (*Piceae glauca*), Lutz spruce (*P. glauca x sitchensis*), and Sitka spruce (*P. sitchensis*), and to a lesser extent, black spruce (*P. mariana*) (USDA Forest Service 2024). As of 2021 the infestation was reported as far north as the northern Matanuska-Susitna Borough/lower Denali Borough to the north and the Chugach National Forest and the Kenai Peninsula to the south. (USDA Forest Service 2024).

Amber-marked birch leaf miner (*Profenusa thomsoni* (Konow)), a non-native insect herbivore of Alaskan birch (*Betula neoalaskana*), was first reported in Anchorage in the 1990s, likely from imported ornamental trees. Since the early 2000s it has spread as far north as Fairbanks, where it heavily infests native birch trees in urban settings and attacks forested areas to a lesser extent (USDA Forest Service 2024). Larval mining of photosynthetic tissue occurs later in the summer, when most tree growth is completed, and long-term effects of *P. thomsoni* on native Alaskan birch health remain unknown.

Tree-level surveys are relatively simple and quick and can add information depth to remote-sensed forest monitoring surveys (Zuidema and van der Sleen 2022). In 2021-2022 we indexed visual signs of tree stress of 359 deciduous and 309 coniferous trees from five sites near Fairbanks, Alaska, as part of a wider study on forest resilience. Sites encompassing different elevations, moisture regimes and burn severities were imaged by NASA aircraft in 2017-2022 using Airborne Visible/Infrared Imaging Spectrometer Next Generation (AVIRIS-NG) hyperspectral technology for the Arctic and Boreal Vulnerability Experiment (ABoVE) (National Aeronautics and Space Administration 2022; 2024). The NASA ABoVE program has collected numerous AVIRIS-NG images in arctic and boreal ecosystems at 5-m resolution (425 spectral bands, 5 nm intervals), providing full hyperspectral reflectance raster data.

We predicted that resilient forests would show little or no visual signs of tree damage from pests/disease/browning, and that vulnerable sites would show moderate to severe signs of tree damage, regardless of species, tree demography, site characteristics or burn history.

## Methods

### Study sites

GT surveys were made of five sites within four AVIRIS scenes (Figure 1) between June and August of 2021 and 2022 at peak leaf condition. 44 polygons inside the AVIRIS scenes representing typical upland and lowland forest (35,000 m^2^ average area) were selected for plant and soil measurements, including tree damage assays. 25 of the polygons were of different burn severities determined by the Alaska Monitoring Trends in Burn Severity index (Nelson 2022). Plots were situated at least 50 m from trails or roads to minimize edge effects. We used a Garmin inReach Explorer (Garmin, Inc., Olathe, USA) supplied by NASA to geolocate sampling plots.

**Figure 1.**
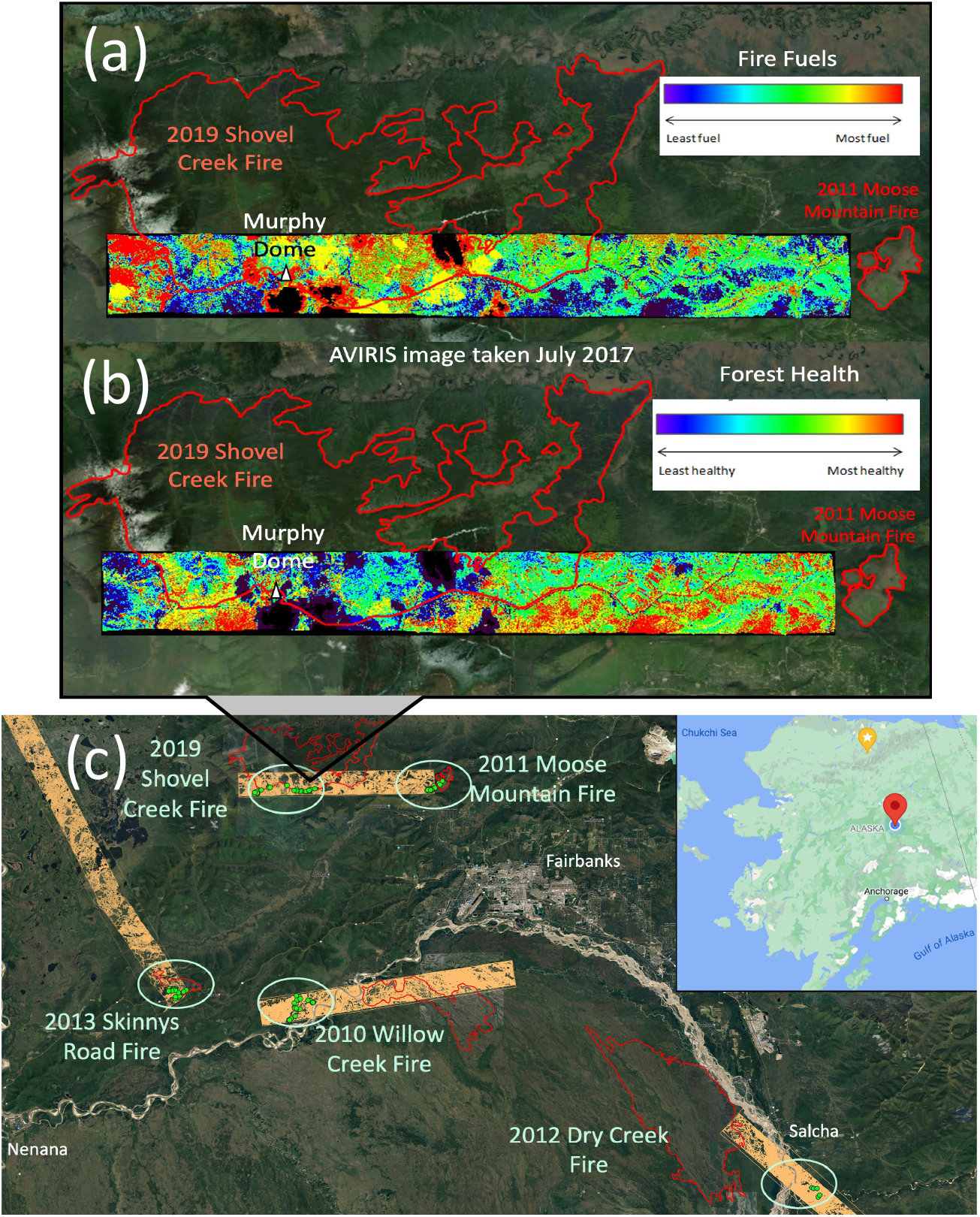
(a-b): Analysis of canopy pigments, water content, and non photosynthetic carbon imaged from 29 hyperspectral bands of an AVIRIS-NG data setof Shovel Creek, Alaska(Murphy Dome is shown in (a-b) for reference).(a) shows high fire fuel loads in 2017; (b) shows declining forest health in2017 relative to the 2019 Shovel Creek Fire scar(red perimeter). (c)Study area consists of 44 ground-truth (GT) plots(green dots inside ovals)across four AVIRIS-NG scenes (orange rectangles) imaged by NASA aircraft in 2017for the Arctic and Boreal Vulner ability Experiment (ABoVE campaign.

### Spectral analysis of forest health and fire fuels

Prior to GT we performed spectral analyses of forest health and fire fuel loads of landscapes imaged in the 2017 AVIRIS scene that includes Shovel Creek, Murphy Dome, and Moose Mountain (Figure 1 a,b). Using ENVI 5.5 software (L3 Harris Geospatial Solutions 2022) we calculated indices of canopy pigment, water content and non-photosynthetic carbon from the initial 29-band subset, which provided background for GT surveys of visual tree damage (Table S1).

### Tree and cover measurements

The 668 focal trees used in the study (359 deciduous, 309 coniferous) formed the dominant forest canopies. These consisted of Alaska paper birch (*Betula neoalaskana*), aspen (*Populus tremuloides*), balsam poplar (*Populus balsamifera*), black spruce (*Picea mariana*) and white spruce (*Picea glauca*). Measurements of 20 randomly selected focal trees plot^-1^ (average: 16 trees plot^-1^) were made inside circular plots (1/30 ha, fixed radius: 10.36m) after Andersen et al. (2011). In post-burned sites where no mature trees were found, we randomly selected recruited seedlings (≤ 1 m height) or saplings (≤ 3 m height) forming the tallest vegetation canopy; at our sites conifer seedlings were found in the undercanopy below the deciduous seedlings and were therefore excluded. Height and diameter at breast height (DBH) of each tree was measured. Using indices of 0 - 5 (0 = no damage/healthy looking trees, 5 = severe damage) we scored each tree for: A) leaf damage from holes, mining, or leaf loss; B) stem damage as changes in stem color, texture, or stem growth; holes or sap ooze; tissue loss caused by herbivory, breakage, pests or pathogens; and heartwood rot assessed from cores extracted with a 5mm increment borer; C) leaf wilt (broadleaf species only); and D) non-photosynthetic portion of the canopy, aka “browning”. Average damage tree^-1^ was calculated as the mean of the four scores (Figure 2). In addition, we characterized the overall crown color tree^-1^ (> 50% of the crown area) as either green, yellow, white, or brown, modified from Boucher and Mead (2006). Details about likely damage sources were identified to species where possible (e.g., browsers, insects, and pathogens).

**Figure 2.**
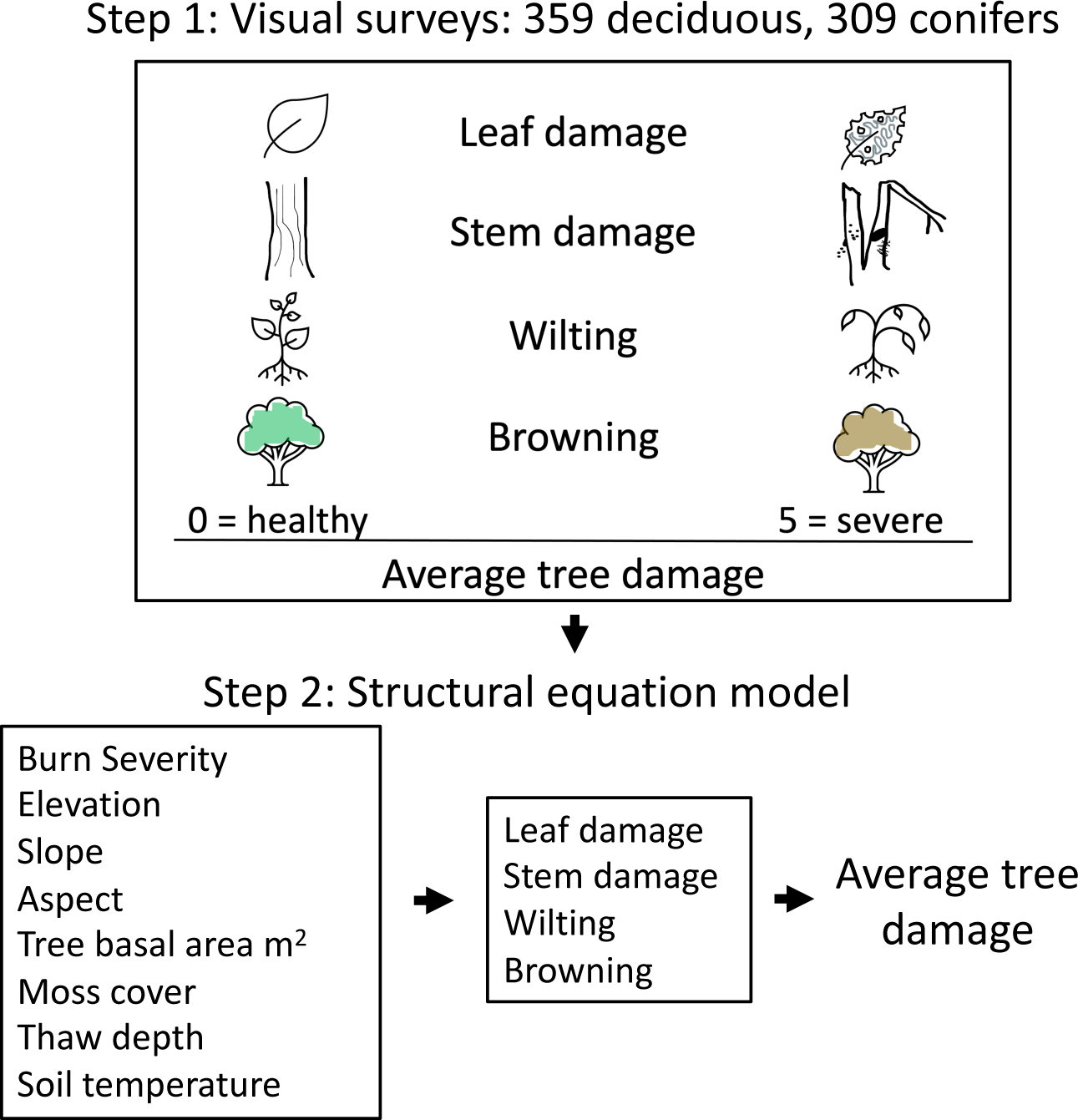
Schematic of visual tree damage assays. Values ranging from 0 (healthy looking trees) to 5 (severe damage) were assigned to 359 deciduous (birch, aspen, and balsam poplar) and 309 coniferous trees (black spruce, white spruce) for four indices of tree damage: leaf damage, holes, mining, tissue loss) stem damage (various; see Methods), wilting (broadleaf species only) and browning (canopy color change including nonphotosynthetic/ necrotic tissue). Average tree damage was calculated as the mean the four indices for each tree.

### Statistical analysis

SEMs (structural equation models) were compared to explain the contribution of environmental variables on leaf damage, stem damage, wilting and browning, and how these four components contributed to average tree damage. We selected eight independent variables describing typical landscape characteristics (*r* < 0.50) (Table S2): 1) MTBS burn severity: 0 = unburned, 1-2 = light burn, 2-3 = moderate burn, 3-4 = severe burn (Nelson 2022); 2) tree basal area m^2^; 3) elevation (in meters); 4) percent slope; 5) aspect (degrees); 6) percent cover of ground moss; 7) maximum thaw depth (in meters); and 8) maximum soil temperature (°C). Model fit converged after 1000 iterations. A reduced SEM model was selected for lowest AICc values (Table S3) and significant regression relationships (*P* ≤ 0.1) (Table S4).

One-way ANOVA of linear models was performed for each damage component across MTBS burn categories (4 levels: unburned, lightly burned, moderately burned, and severely burned); by species (5 levels; see above for species details), site, age class (two levels: seedling/sapling, or tree), elevation class (3 levels representing lowlands, mid-elevation, and uplands), and stem size classes (10 levels, derived from DBH). Interactive terms were precluded by the large number of categories.

To understand mechanisms of damage, we calculated the percentage of total trees sorted by species (conifer, deciduous) or primary canopy color (brown, green, white, yellow) experiencing the damage components described above across 15 likely sources of damage we identified in surveys relative to undamaged trees. Damage categories included mammalian browsers, insects, pathogens, leaf chlorosis, stem breakage, sap ooze, and undercanopy browning. In our survey we found only leaf defoliating insects. For pathogens we combined leaf and stem pathogens (e.g., cankers and heartwood rot) into a single category. Analysis was done in R (R Core Team 2023) and JMP Pro 16 (JMP Statistical Discovery 2023).

## Results

Spectral analyses of the 2017 AVIRIS-NG scene encompassing Shovel Creek, Murphy Dome, and Moose Mountain revealed higher indices of fire fuel loads and declining forest health seen as changes in canopy pigment, reduced canopy water content, and non-photosynthetic carbon (Figure 1a-b).

Average tree damage from best SEM models was explained, in descending order of model variance, by stem damage, wilting, leaf damage, and browning (Figure 3, Table S4). Stem damage was largely explained by post-burned, sloping sites, E and S facing sites, sites with less moss cover and warmer soil temperatures, and to some degree by sites at higher elevations (Figure 3, Table S4). Wilting was associated with severely burned sites with less moss cover, lower elevation sites, NW facing sites with colder soils and shallower thaw depths, and to a lesser extent, early successional sites with abundant seedlings and saplings (Figure 3, Table S4). Higher average leaf damage was found in sites at lower elevations, in E and S facing, steeply sloped sites, and in sites with warmer, shallower soils and less moss cover (Figure 3, Table S4). Browning was explained by sites featuring smaller trees with shallower, colder soils with more moss cover characteristic of black spruce sites, which, to some degree, were found at lower elevations (Figure 3, Table S4).

**Figure 3.**
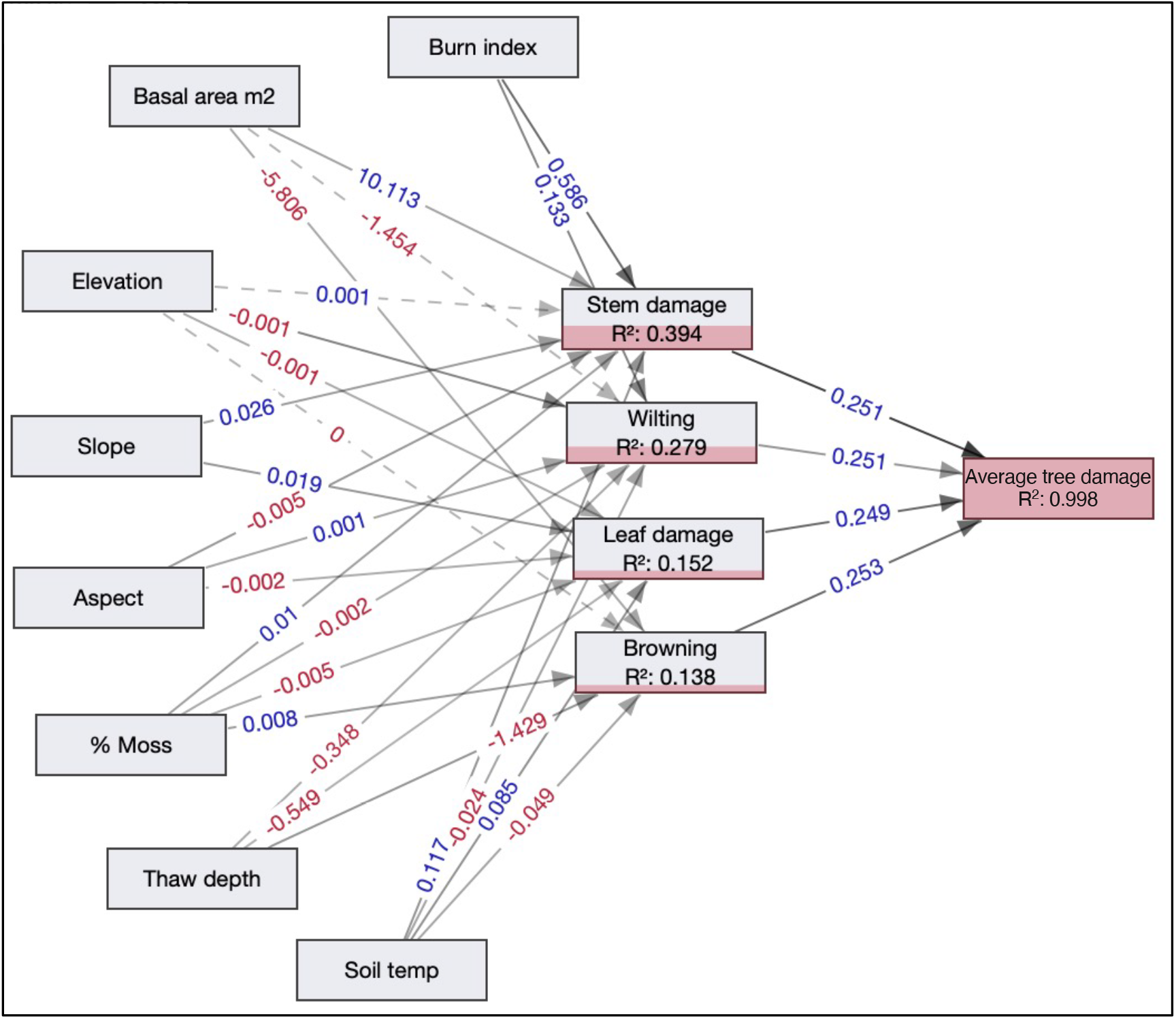
Structural equation model shows regression relationships of eight environmental characteristics on tree damage components of 359 deciduous and 309 conifer trees near Fairbanks, Alaska. Numbers along arrows show change from the mean. Red numbers indicate negative relationships; blue numbers indicate positive relationships. Solid line shows statistically significant relationships (*P* < .05), dashed line shows marginally significant relationships (*P* < .1). Red shading indicates proportional variance explained. Models converged after 1000 iterations.

Burn severity was a highly significant variable, explaining over 15% of variance in average tree damage including over 23% of stem damage in one-way ANOVA of linear models (Table 1). Across burn severities, average tree damage was light to moderate in severely and lightly burned sites and low elsewhere (Figure 4a). These averages were higher due to moderately severe stem damage seen in recovering severely burned sites, and moderate leaf damage, browning, and wilting in lightly burned sites (Figure 4a; Table 1). By contrast, trees in moderately burned and unburned sites showed low average damage (Figure 4a).

**Table 1.**
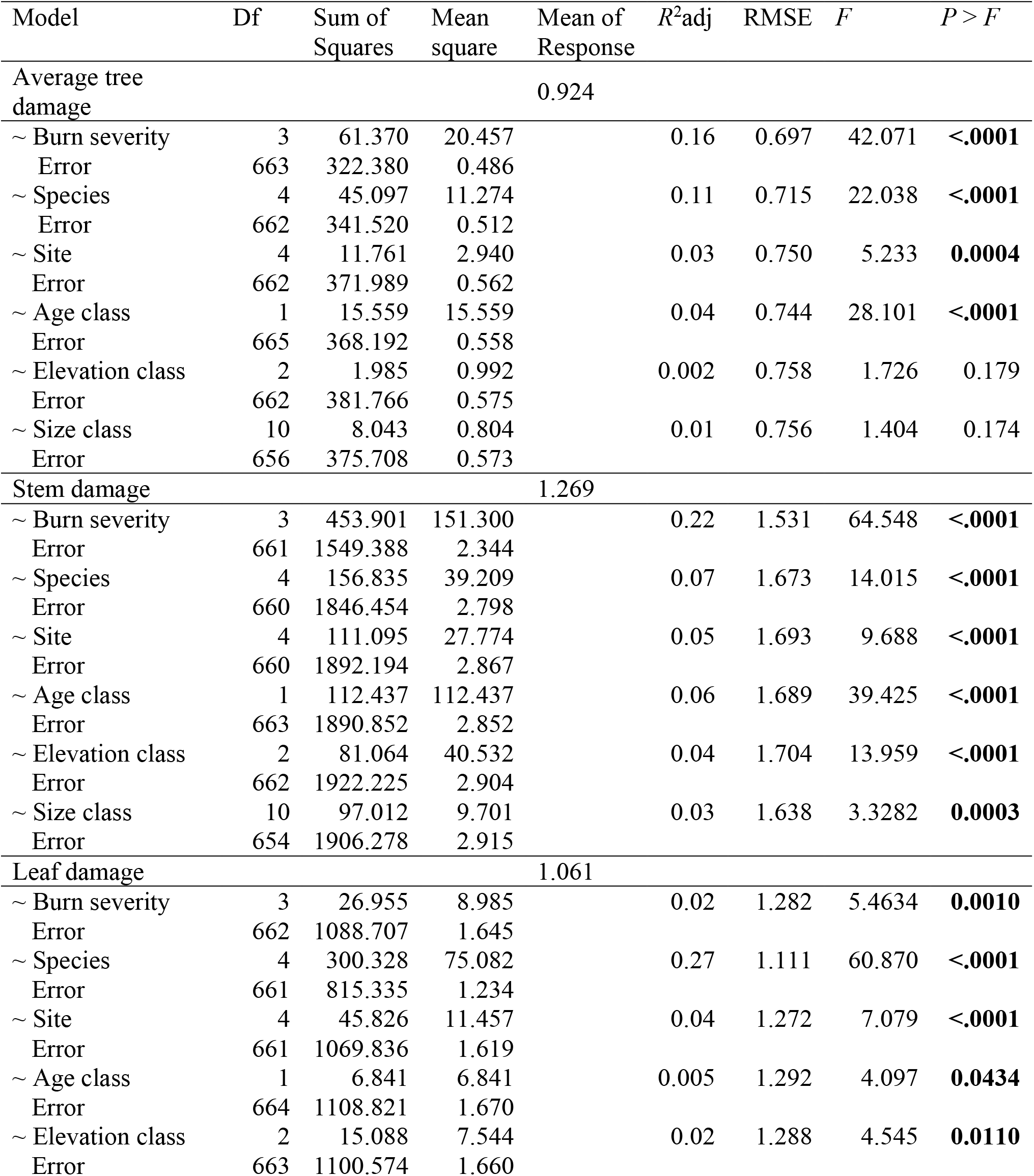
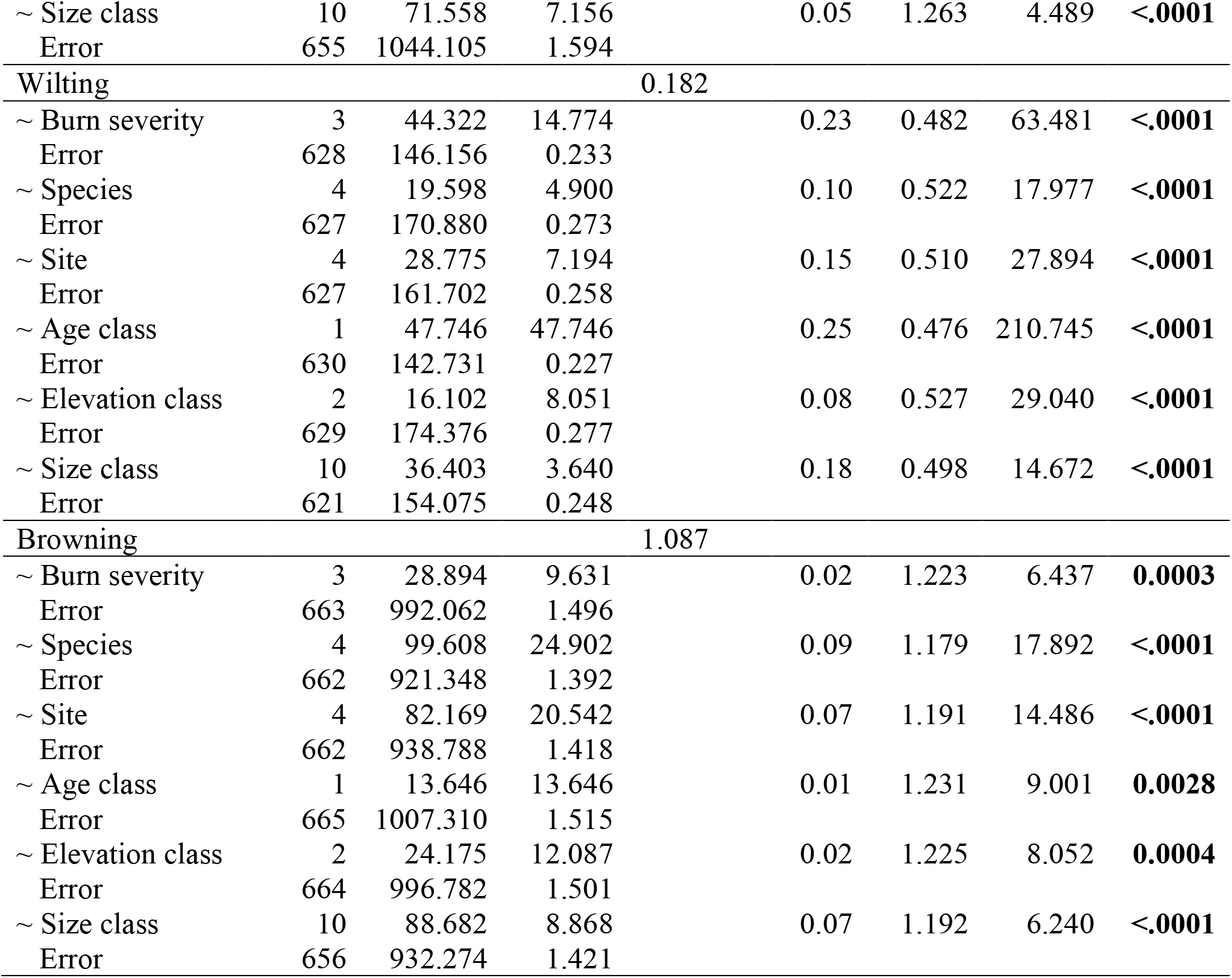
One-way Analysis of Variance of tree damage (Average tree damage, stem damage, leaf damage, wilting, and browning) by burn severity, tree species, site, age class (tree versus seedling), elevation class, and stem diameter at breast height (DBH) size classes, based on visual assays of 668 trees observed in AVIRIS-NG imaged forests near Fairbanks, Alaska. Bold numbers indicate model significance at *P* < 0.05.

**Figure 4.**
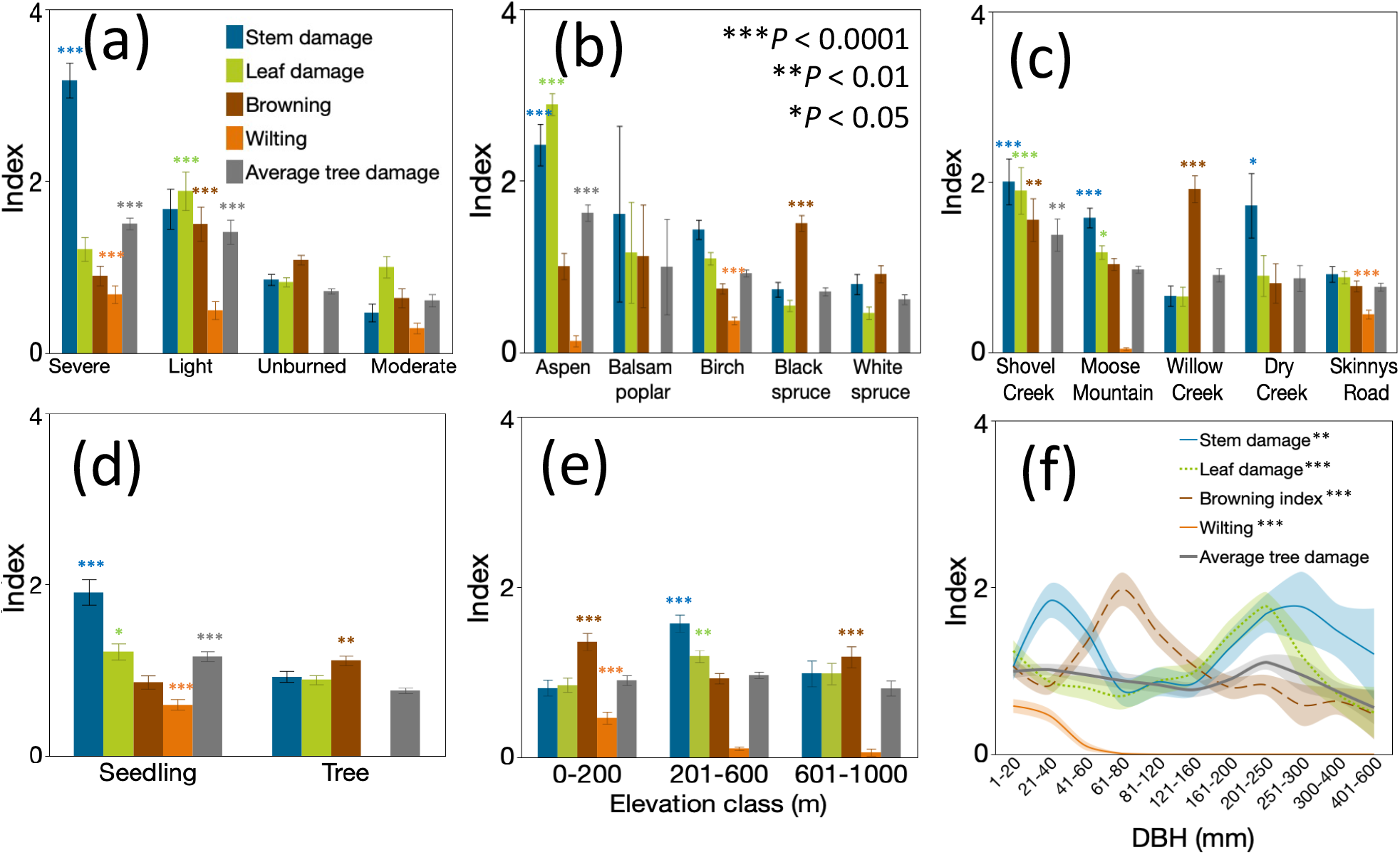
Tree damage index means from visual surveys of 668 trees at five sites near Fairbanks, Alaska by (a) MTBS burn severity classes; (b) species; (c) site; (d) age class (seedling/sapling or tree); (e) elevation categories comprising lowlands, mid-elevation, and uplands; and (f) diameter at breast height (DBH) size classes. Index values in A-D are shown in descending order by average tree damage. Error bars in (a-e) and shaded bands in (f) show 1 standard error. Stars indicate model significance in one-way analysis of variance.

Tree damage varied significantly within and across species (Figure 4b). Species effects explained over 11% of the variance in average tree damage, including 26% of the variance in leaf damage in one-way ANOVA (Table 1). Aspen trees showed moderately high damage scores, mainly explained by leaf and stem damage that was three to six times higher than for other species (Figure 4b). Black spruce damage was mainly explained by moderate browning (Figure 4b). Birch showed the highest wilting index across species, although average birch wilting was low (Figure 4b).

Sites showed significant variation in tree damage, although site effects only explained 2.5% of the variance of average tree damage in one-way ANOVA (Table 1). Shovel Creek showed the highest average tree damage, which was mainly explained by moderate leaf and stem damage and moderate browning (Figure 4c). Moose Mountain and Dry Creek also showed moderate stem damage, while Willow Creek showed moderate browning, mainly associated with conifers (Figure 4b, c). Wilting, low overall, was highest at Skinny’s Road (Figure 4c).

Seedlings suffered higher average tree damage than mature trees, mainly from moderate stem damage and low to moderate leaf damage and wilting (Figure 4d). Mature trees showed the most browning, however values were below moderate (Figure 4d).

Elevation classes representing lowland, mid elevation, and upland sites, and stem diameter classes were insignificant for average tree damage in one-way ANOVA of linear models (Table 1), because damage means did not differ across elevational or stem diameter gradients (Figure 4e, f). However, both variables had significant effects on the four components of average tree damage in one-way ANOVA of linear models (Table 1). Elevation accounted for 8% and 4% of the variance in browning and stem damage, respectively (Table 1). Lowlands and uplands showed the highest browning scores, although these values were low overall, while middle elevations showed significant moderate stem damage and low to moderate leaf damage (Figure 4e).

Stem diameter at breast height (DBH) classes explained 7% and 18% of the variance in wilting and browning, respectively, in one-way ANOVA of linear models (Table 1). Stem damage was greatest in trees with DBH between 20-40 mm and 250-300 mm, where it reached moderate levels, but not for trees between 60-200 mm (Figure 4f). On average, moderate leaf damage occurred in trees with DBH of 150-300 mm, while moderate browning occurred mainly in smaller trees with DBH between 60-100 mm (Figure 4f). Wilting was primarily confined to the smallest size classes and dropped off once trees reached DBH beyond 60 mm (Figure 4f).

On average, a third of the trees showing insect damage in our survey were deciduous species (Figure 5a). 18-30% of trees showing stem damage, wilt, and canopy browning also showed insect attack (Figure 5b-e). Stem damage by browsers was seen on 22% of trees, mainly deciduous species (Figure 5b), however, damage by a combination of browsers and insects was low (Figure 5). Nearly half of all trees showing leaf damage by insect herbivores were deciduous species (Figure 5c). By contrast, damage from a combination of insects and pathogens was seen on more than 10% of surveyed trees, mainly deciduous species (Figure 5a), and was associated with higher proportions of stem damage and wilting (Figure 5b, d). Pathogens (cankers, leaf spots, blights, heartwood rot) were found in nearly 8% of trees across species (Figure 5a-e). For trees with stem damage, 12% showed signs of pathogen attack (Figure 5b). Nearly 40% of coniferous trees showed some form of undercanopy browning (Figure 5e, j), and in post-burned sites, live conifers had scorched stems (Figure 5b). The number of trees suffering no visible signs of damage was low, around 5% (Figure 5a, f), mostly explained by a relatively high proportion (17%) of trees across species showing no signs of wilting (Figure 5d, i). Across damage components and likely damage sources, including leaf damage, tree canopies were primarily green (Figure 5f-j). The exceptions were trees with chlorotic leaves and trees with undercanopy browning, although smaller numbers of trees scored for sap ooze and stem breakage + pathogens had browner canopies (Figure 5f-j).

**Figure 5.**
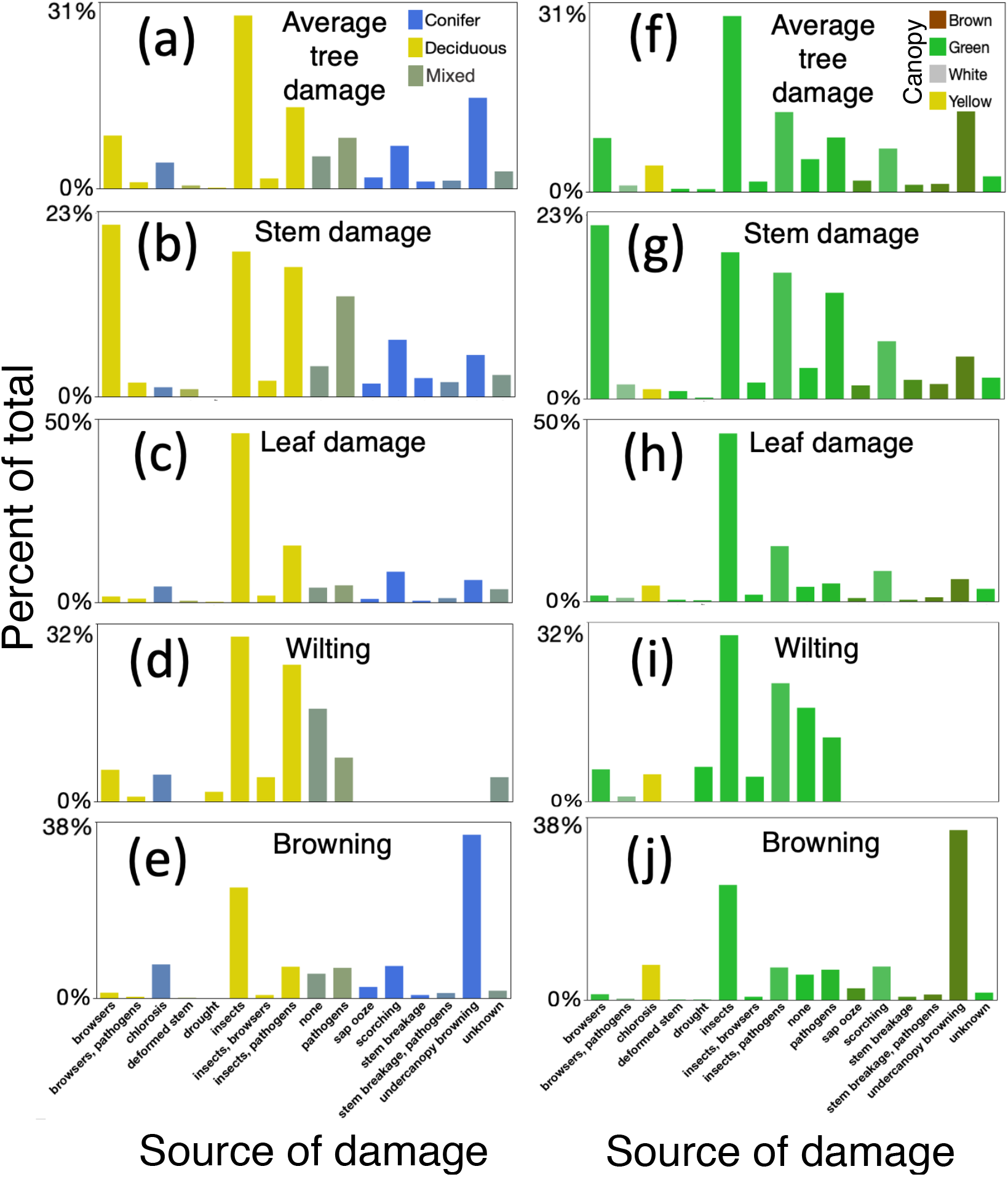
Percent of 668 live trees from five forest sites near Fairbanks, Alaska scored for four components of average tree damage from 0 - 5 (none to severe) across 16 likely damage sources determined in visual surveys. (a-e) shows proportions by forest type conifer (yellow bars), deciduous (blue) and mixed (grey-green); (f-j) shows the same information by dominant canopy color tree-1: brown, green, white and yellow (darker or lighter colors show mixtures). The *x*-axis shows 15 likely sources of damage determined from visual surveys; “none” shows the proportion of undamaged trees. “Insects” include foliar insect species only; “pathogens” include leaf and stem pathogens;“unknown” is unidentified sources of damage.

## Discussion

Our tree-level damage survey of 668 focal trees in forests near Fairbanks, Alaska found low to moderately low tree damage overall, however, damage to leaves and stems, the severity of browning and wilting, and likely sources of damage were varied and complex. Damage scores appeared to be significantly influenced by burn severity, species, tree age and size, location, and abiotic characteristics. Sources of damage used in our proportional comparisons (Figure 5) were derived from 25 different types of observations. The most obvious signs of damage were found in the leaves and stems of deciduous trees (Figure 6 a-g). Leaf damage was mainly caused by larval stages of various insect herbivores, and to a lesser extent by pathogens (Figure 6 a-d). Stem damage was mainly caused by mammalian herbivores and stem breakage and rot of (Figure 6g-i).

**Figure 6.**
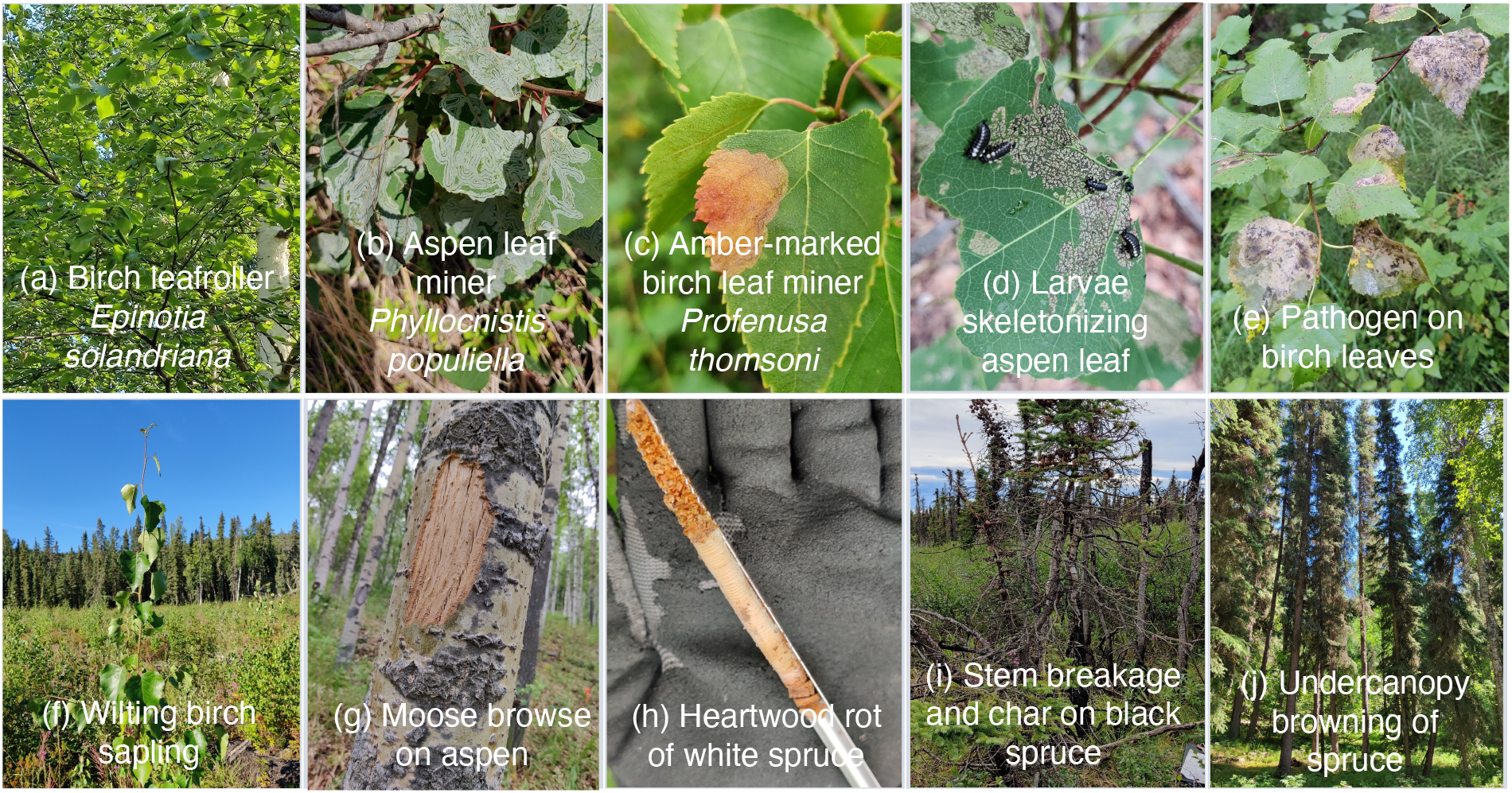
Visual documentation of damage includes (a) birch tree infested with leafroller, Bonanza Creek Experimental Forest; (b) aspen leaf miner (*Phyllocnistis populiella*), Moose Mountain burn perimeter; (c) amber-marked birch leaf miner (*Profenusa thomsoni*), Skinny’s Road burn perimeter; (d) larvae skeletonizing aspen leaf, Moose Mountain Ski area; (e) Leaf pathogens attacking birch leaves in unburned forest near Dry Creek burn perimeter, Salcha, Alaska; (f) wilting birch sapling, Skinnys Road burn perimeter; (g) moose herbivory of aspen bark, Moose Mountain Ski area; (h) heartwood rot of white spruce in unburned forest near Dry Creek burn perimeter, Salcha, Alaska; (i) stem breakage and char on live black spruce, Shovel Creek burn perimeter; and (j) undercanopy browning of white spruce, unburned conifer forest near Skinny’s Road burn perimeter.

Although over half of aspen tree canopies surveyed were green, our results suggest that aspen in the study area is most vulnerable to insect attack, mainly aspen leaf miner (*P. populiel*), in agreement with other research (Wagner and Doak 2013). However, infested aspen in the Alaskan interior have not shown a strong relationship to NDVI changes (Boyd et al. 2021). Other aspen damage included stem browse by mammals and heartwood rot. The aspen we surveyed included mature trees on unburned upland ski slopes (Moose Mountain) and seedling and sapling recruits in burn scars. At our sites aspen forest occurred on the steepest slopes at mid to upland elevations (300-500 m). These plots were associated with warmer, drier, shallower soils with less moss cover typical of well-drained early to mid-successional sites.

Remote sensing has detected NDVI changes in deciduous woody shrubs related to severe insect attack and recovery at northern latitudes (Prendin et al. 2020), however, remote-sensed measurements may not capture true variation in canopy conditions related to insect attack if trees of different heights are attacked (Boyd et al. 2021). Recent research suggests that taller aspen trees in Alaska’s Interior have a greater probability of being attacked by aspen leaf miner due to earlier leaf-out providing optimal windows for oviposition (Tundo 2021).

Damage to conifers was mainly found in stems from pathogens leading to heartwood rot (Figure 6h), mechanical breakage likely from storms and snow loads, and scorching in lightly burned sites (Figure 6i). The most common type of damage seen in conifers, however, was undercanopy browning of mature spruce (Figure 6j). This is likely a typical characteristic of conifer aging caused by self-shading and senescence of lower branches, although it was prominent in small-diameter conifers. Significantly, we did not detect signs of spruce beetle infestation at our sites.

Spruce beetle infestation is linked to warmer temperatures promoting accelerated larval development; however, forest type likely influences response trajectories following beetle outbreaks. In Alaska’s Cook Inlet, high mortality of white spruce in the 1970s led to birch dominated forest (Werner et al. 2006). In the Kenai Peninsula, white spruce mortality from an outbreak in the 1980s led to increased fuel loads from dead trees and growth of flammable grasses (Werner et al. 2006). In the 1990s, beetle infestations in the Kenai Peninsula did not alter forest regeneration trajectories in lowland mixed forest, demonstrating resilience (Boucher and Mead 2006). Alaska’s Interior is characterized by large stands of black spruce, which is less attacked by spruce beetle than white spruce. However, as spruce beetle moves north into the Alaskan Interior with climate warming, conifer forest may be increasingly vulnerable to severe fires due to increased fuel loads of beetle-killed white spruce in proximity to the more flammable black spruce stands (Hansen et al. 2016). Increasing fire frequency and severity is a driver of forest change in Alaska’s Interior from black spruce to deciduous forest (Mann et al. 2012).

*P. thomsoni* is a relatively recent pest in Alaska, and its effects on Interior birch forests remain largely understudied. In the summer of 2022, we found light to moderate infestations on mature forest trees and saplings, and heavy infestations on mature trees in the city of Fairbanks (Figure S1). Previous long term biological control studies found low growth of recovered Alaskan trees following a 2008 infestation (Andersen et al. 2021; Van Driesche et al. 2023). One possible explanation was invasion of another leaf-mining species, *Heterarthrus nemoratus* (Fallén). Although there has been no observed tree mortality of birch following *P. thomsoni* infestations, recent research suggests infestations of novel pests may lead to the decline of some Alaskan tree species over time (Ruess et al. 2021).

## Conclusion

Our study used a simple and fast GT method to visually assess signs of tree damage in NASA ABoVE remote-sensed forests near Fairbanks, Alaska. In particular, we found tree-level analysis useful following spectral analysis of a 2017 AVIRIS-NG dataset revealing high forest stress and fuel loads in the Shovel Creek watershed which burned in 2019. Our survey sheds light on the importance of forest stress studies integrated at different spatial scales, particularly since we found severe infestations of the amber-marked birch leaf miner, of a relatively new forest pest in Alaska, that is now infesting birch trees in the Interior.

## Author contributions

DCH and CSP conceived of the ideas and designed the methodology; DCH conducted field sampling; DCH analyzed the data, with CSP and others. All authors contributed critically to the drafts and gave their final approval.

## Acknowledgments

We thank the Dena People of Interior Alaska for access to the sites where we conducted our research. This research was sponsored by the National Aeronautics and Space Administration ®NASA© 2024 through a contract with Oak Ridge Associated Universities, and NASA Ames Research Center (STRIVES 20220017558). The views and conclusions contained in this document are those of the authors and should not be interpreted as representing the official policies either expressed or implied of NASA or the U.S. Government. The U.S. Government is authorized to reproduce and distribute reprints for government purposes notwithstanding any copyright notation herein. Special thanks to Emily Reast and Lesley Tilghman and for assistance in the field, and to the University of Alaska Fairbanks for logistical support.

