## Supplemental Files for "Comparisons of Tree Damage Indicators in Five NASA ABoVE Forest Sites Near Fairbanks, Alaska"

**Supplemental Information**

**Table S1**. Indices of forest health and fire fuels used in ENVI 5.5 to classify AVIRIS imagery of Moose Mountain/Shovel Creek area near Fairbanks, AK, from a subset of 29 hyperspectral bands (see Figure 1).

| **Forest Health Indices** | | | |
|---|---|---|---|
| **MRENDVI (p750-p705/p750+p705-2*p445** | **Leaf Pigment (Cretinoid Reflectance Index)** | **Canopy water content** | |
| 0.2 | Anthocyanin Reflectance Index 2 (ARI2) = p800[1/p550 - 1/p700] | Normalized Difference Water Index (NDWI = (p857-p1241)/(p857+p1241) | |
| **Fire Fuels Indices** | | | |
| **MRENDVI (p750-p705/p750+p705-2*p445** | | **Canopy water content** | **Dry or senescent carbon index** |
| 0.2 | | Normalized Difference Water Index (NDWI = (p857-p1241)/(p857+p1241) | Cellulose Absorption Index (CAI) = 0.5(P2000+P22000)-P2100 |

**Table S2**. Correlation matrix of explanatory variables used in Structural Equation Models of tree damage.

| Row | Burn index | Basal area m2 | Elevation | Slope | Aspect | % Moss | Thaw depth | Soil temp |
|---|---|---|---|---|---|---|---|---|
| Burn index | 1 | -0.347 | -0.051 | -0.126 | -0.021 | -0.364 | 0.061 | 0.176 |
| Basal area m2 | -0.347 | 1 | -0.219 | 0.041 | 0.115 | 0.130 | 0.100 | 0.121 |
| Elevation | -0.051 | -0.219 | 1 | 0.283 | 0.000 | -0.061 | -0.016 | 0.142 |
| Slope | -0.126 | 0.041 | 0.283 | 1 | 0.073 | -0.241 | -0.200 | 0.416 |
| Aspect | -0.021 | 0.115 | 0.000 | 0.073 | 1 | 0.226 | 0.111 | 0.216 |
| % Moss | -0.364 | 0.130 | -0.061 | -0.241 | 0.226 | 1 | -0.161 | -0.306 |
| Thaw depth | 0.061 | 0.100 | -0.016 | -0.200 | 0.111 | -0.161 | 1 | -0.130 |
| Soil temp | 0.176 | 0.121 | 0.142 | 0.416 | 0.216 | -0.306 | -0.130 | 1 |

**Table S3.** Model selection of Structural Equation models. Model fit converged after 1000 iterations. A reduced SEM model (Model 15) was selected as the best model for low AICc values.

| Model Name | -2 Log Likelihood | No. of Parameters | AICc | AICc Weight | BIC | ChiSquare | DF | Prob> ChiSq | CFI | RMSEA | Lower 90% | Upper 90% |
|---|---|---|---|---|---|---|---|---|---|---|---|---|
| Unrestricted | 35207.3703 | 104 | 35454.1624 | 1 | 35883.8162 | 0 | 0 | | 1 | 0 | 0 | 0 |
| Independence | 41242.1525 | 26 | 41296.3429 | 0 | 41411.264 | 6034.7823 | 78 | <.0001 | 0 | 0.3381 | 0.3309 | 0.3454 |
| Model 1 | 36273.7632 | 61 | 36408.2451 | 0 | 36670.5248 | 1066.393 | 43 | <.0001 | 0.8282 | 0.1888 | 0.179 | 0.1986 |
| Model 2 | 36273.7696 | 60 | 36405.8289 | 0 | 36664.0269 | 1066.3994 | 44 | <.0001 | 0.8284 | 0.1865 | 0.1769 | 0.1963 |
| Model 3 | 36273.8417 | 59 | 36403.4864 | 0 | 36657.5947 | 1066.4714 | 45 | <.0001 | 0.8285 | 0.1843 | 0.1748 | 0.194 |
| Model 4 | 36274.4794 | 58 | 36401.7175 | 0 | 36651.7281 | 1067.1092 | 46 | <.0001 | 0.8286 | 0.1823 | 0.1729 | 0.1919 |
| Model 5 | 36275.7784 | 57 | 36400.6177 | 0 | 36646.5228 | 1068.4081 | 47 | <.0001 | 0.8285 | 0.1804 | 0.1711 | 0.1898 |
| Model 6 | 36277.5761 | 56 | 36400.0246 | 0 | 36641.8163 | 1070.2059 | 48 | <.0001 | 0.8284 | 0.1786 | 0.1693 | 0.1879 |
| Model 7 | 36279.6551 | 55 | 36399.7205 | 0 | 36637.391 | 1072.2849 | 49 | <.0001 | 0.8282 | 0.1768 | 0.1677 | 0.1861 |
| Model 8 | 36282.9276 | 54 | 36400.6177 | 0 | 36634.1592 | 1075.5574 | 50 | <.0001 | 0.8278 | 0.1752 | 0.1662 | 0.1844 |
| Model 9 | 36279.6551 | 55 | 36399.7205 | 0 | 36637.391 | 1072.2849 | 49 | <.0001 | 0.8282 | 0.1768 | 0.1677 | 0.1861 |
| Model 10 | 36282.8676 | 54 | 36400.5576 | 0 | 36634.0992 | 1075.4973 | 50 | <.0001 | 0.8278 | 0.1752 | 0.1662 | 0.1844 |
| Model 11 | 36286.5173 | 53 | 36401.8397 | 0 | 36631.2445 | 1079.147 | 51 | <.0001 | 0.8274 | 0.1737 | 0.1648 | 0.1828 |
| Model 12 | 36282.8676 | 54 | 36400.5576 | 0 | 36634.0992 | 1075.4973 | 50 | <.0001 | 0.8278 | 0.1752 | 0.1662 | 0.1844 |
| Model 13 | 36289.7904 | 52 | 36402.753 | 0 | 36628.0134 | 1082.4202 | 52 | <.0001 | 0.827 | 0.1722 | 0.1634 | 0.1812 |
| Model 14 | 36282.8676 | 54 | 36400.5576 | 0 | 36634.0992 | 1075.4973 | 50 | <.0001 | 0.8278 | 0.1752 | 0.1662 | 0.1844 |
| Model 15 | 36279.6551 | 55 | 36399.7205 | 0 | 36637.391 | 1072.2849 | 49 | <.0001 | 0.8282 | 0.1768 | 0.1677 | 0.1861 |

**Table S4.** Regression goodness of fit table used in the best SEM model (Model 15). Model regressed eight environmental variables (burn index, elevation, slope, aspect, percent cover of moss, maximum summer thaw depth, maximum summer soil temperature, basal area of tree) on four components of average tree damage: leaf damage, stem damage, browning, and wilting (each was given a score from 0 to 5 for 708 focal trees). Table shows estimates from the mean of each regression, standard error, Wald Z-statistic, and *P*-values.

| **Regressions** | **Estimate** | **Std Error** | **Wald Z** | **Prob>\|Z\|** |
|---|---|---|---|---|
| Stem damage → Average tree damage | 0.2514 | 0.0006 | 398.4399 | 0.0000 |
| Leaf damage → Average tree damage | 0.2485 | 0.0009 | 276.4643 | 0.0000 |
| Wilting → Average tree damage | 0.2511 | 0.0019 | 128.9582 | 0.0000 |
| Browning → Average tree damage | 0.2527 | 0.0009 | 279.2773 | 0.0000 |
| Burn index → Stem damage | 0.5859 | 0.0477 | 12.2868 | 0.0000 |
| Basal area m2 → Stem damage | 10.1127 | 2.2770 | 4.4412 | 0.0000 |
| Slope → Stem damage | 0.0255 | 0.0048 | 5.3049 | 0.0000 |
| Aspect → Stem damage | -0.0045 | 0.0007 | -6.4574 | 0.0000 |
| % Moss → Stem damage | 0.0103 | 0.0021 | 4.8242 | 0.0000 |
| Soil temp → Stem damage | 0.1168 | 0.0187 | 6.2393 | 0.0000 |
| Elevation → Leaf damage | -0.0007 | 0.0002 | -3.0586 | 0.0022 |
| Slope → Leaf damage | 0.0194 | 0.0038 | 5.0843 | 0.0000 |
| Aspect → Leaf damage | -0.0015 | 0.0006 | -2.6678 | 0.0076 |
| % Moss → Leaf damage | -0.0047 | 0.0017 | -2.8374 | 0.0045 |
| Thaw depth → Leaf damage | -0.5488 | 0.2446 | -2.2438 | 0.0248 |
| Soil temp → Leaf damage | 0.0850 | 0.0148 | 5.7562 | 0.0000 |
| Burn index → Wilting | 0.1330 | 0.0161 | 8.2691 | 0.0000 |
| Elevation → Wilting | -0.0009 | 0.0001 | -9.1481 | 0.0000 |
| Aspect → Wilting | 0.0010 | 0.0002 | 3.9073 | 0.0001 |
| % Moss → Wilting | -0.0023 | 0.0007 | -3.0399 | 0.0024 |
| Thaw depth → Wilting | -0.3482 | 0.1050 | -3.3174 | 0.0009 |
| Soil temp → Wilting | -0.0236 | 0.0064 | -3.7134 | 0.0002 |
| Basal area m2 → Browning | -5.8056 | 1.6796 | -3.4565 | 0.0005 |
| % Moss → Browning | 0.0077 | 0.0015 | 5.0428 | 0.0000 |
| Thaw depth → Browning | -1.4294 | 0.2303 | -6.2072 | 0.0000 |
| Soil temp → Browning | -0.0487 | 0.0134 | -3.6388 | 0.0003 |
| Elevation → Stem damage | 0.0005 | 0.0003 | 1.9127 | 0.0558 |
| Elevation → Browning | -0.0004 | 0.0002 | -1.8112 | 0.0701 |
| Basal area m2 → Wilting | -1.4541 | 0.8103 | -1.7945 | 0.0727 |

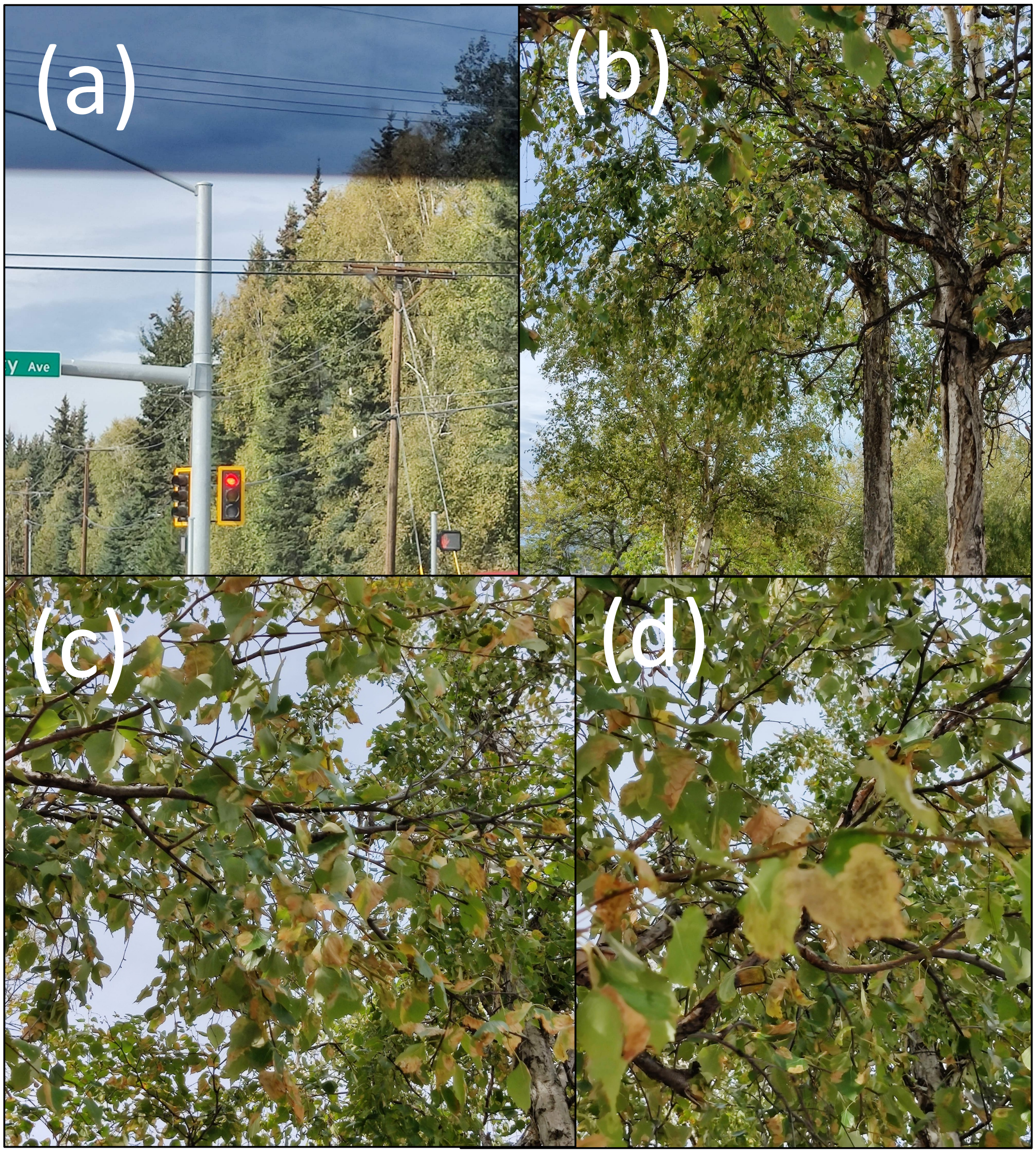


**Figure S1.** Amber-marked birch leaf miner damage (*Profenusa thomsoni* (Konow)) causing browning on mature birch trees in Fairbanks, Alaska on (a) University Avenue and Geist Road; and (b-d) 7th Avenue. Photographed by the author August 22, 2022.
